# Horizontal acquisition of nicotine catabolism gene cluster drives the assembly of tobacco root microbiota community

**DOI:** 10.64898/2025.12.24.696441

**Authors:** Tomohisa Shimasaki, Yui Nose, Sachiko Masuda, Arisa Shibata, Tsubasa Shoji, Shuhei Yabe, Maiko Furubayashi, Yoshitomo Kikuchi, Ken Shirasu, Kazufumi Yazaki, Yasunori Ichihashi, Akifumi Sugiyama, Ryohei Thomas Nakano

**Author notes:** **Corresponding author:** Tomohisa Shimasaki, Department of Biological Sciences, Faculty of Science, Hokkaido University, Sapporo, Hokkaido 060-0810, Japan., Ryohei Thomas Nakano, Department of Biological Sciences, Faculty of Science, Hokkaido University, Sapporo, Hokkaido 060-0810, Japan.

## Abstract

**Background:** Plant roots are hotspots for interactions with soil microbes, where a characteristic bacterial community structure is formed. Plant specialized metabolites often play pivotal roles in this assembly process. However, the molecular basis underlying root microbiota responses to these bioactive compounds, and how such metabolic interactions shape the assembly of host-specific root microbiota, remain largely unknown. Nicotine is a toxic alkaloid predominantly produced by the genus *Nicotiana*, and the genus *Arthrobacter* is known as one of the nicotine-degrading bacteria in the tobacco root microbiota. In this study, we used the tobacco–*Arthrobacter* interaction system as a model and integrated comparative genomics and experimental genetic manipulation assays to uncover the role of bacterial catabolism capacity for host specialized metabolites in shaping host-specific root microbiota.

**Results:** Nicotine catabolism genes are uniquely found in the *Arthrobacter* strains derived from nicotine-containing environments, and this restricted gene distribution is driven by a plasmid-mediated horizontal gene transfer. To assess the ecological consequences of this genomic adaptation in *Arthrobacter* fitness in tobacco roots, we conducted adaptation assays under both *in vitro* and *in planta* conditions using genetically manipulated *Arthrobacter* and tobacco mutants, which are impaired in nicotine catabolism and biosynthesis, respectively. Nicotine improves *Arthrobacter* colonization to the tobacco roots through both catabolism-dependent and catabolism-independent mechanisms. Bacterial community analysis using a synthetic community approach further demonstrated that these metabolic interactions, mediated by tobacco nicotine biosynthesis and its catabolism by *Arthrobacter*, jointly affect root microbiota composition.

**Conclusions:** Our findings illustrated that bacterial catabolic capacity toward host-derived plant specialized metabolites is key for successful root colonization. This metabolic adaptation is driven by plasmid-mediated horizontal gene transfer and ultimately shapes the structure of the overall root microbiota community.

## Background

Plant roots interact with diverse soil microorganisms that form a structured community known as the root microbiota. These root-associated microbes significantly affect plant growth and health by producing plant hormones, improving nutrient uptake, and activating host immunity [1, 2]. Root microbiota profiles vary among plant species at lower taxonomic ranks [3–5], although they eventually converge at the higher levels, such as the order [6–8]. A two-step selection model was hypothesized to explain the assembly process of this characteristic bacterial community structure in the roots [9]. In this model, rhizodeposits serve as initial substrates that shift the soil microbiome community characterized by a core plant microbiome, which is then fine-tuned at lower taxonomic levels by host genotype factors. Accordingly, root microbiota harbor characteristic genomic features that enable adaptation to host-specific root environments, and that such genetic adaptation may underlie the assembly of plant species-specific root microbiota. Indeed, commensal bacteria strains isolated from *Arabidopsis thaliana* or *Lotus japonicus* roots showed a preferential colonization to their native host rather than non-native host, although the taxonomic compositions of both synthetic communities (SynComs) were identical [5].

Root exudates play fundamental roles in assembling the root microbiota. Plant roots secrete up to 25% of their photosynthates into the rhizosphere as root exudates, which contain primary metabolites (such as sugars, amino acids, and organic acids) as well as plant specialized metabolites (PSMs) [10–13]. These metabolites recruit or repel soil microbes, resulting in the formation of characteristic bacterial communities in the roots. While primary metabolites are generally conserved across the plant kingdom, PSMs are produced in a lineage-specific manner and thus are therefore assumed to contribute to plant species-specific root microbiota assembly. In *Arabidopsis thaliana,* for instance, biosynthetic mutants impaired in coumarins [14, 15], triterpenes [16], or tryptophan-derived metabolite pathways [17, 18], harbor root microbiota distinct from those of wild-type plants. Similar modulatory effects have been shown for PSMs such as tomatine in tomato (*Solanum lycopersicum*) [19], triterpene in cucumber (*Cucumis sativus*) [20], and benzoxazinoids (BXs) and flavonoids in maize (*Zea mays*) [21, 22]. Although comparisons between wild-type and biosynthetic mutants using culture-independent amplicon sequencing have well described the impact of PSMs on root microbiota communities, the bacterial genetic basis underlying community shifts driven by PSMs secretion in the root microbiota remain largely unknown.

We previously showed that the nicotine-catabolism genes of the genus *Arthrobacter* are associated with the preferential colonization of tobacco (*Nicotiana tabacum*) roots [23]. Nicotine, a toxic alkaloid predominantly found in *Nicotiana* species, contributes to defense against insect herbivores [24, 25]. Although most of the nicotine synthesized in roots are subsequently translocated to the leaves [26, 27], a portion is also secreted into the rhizosphere [28]. We found that the genus *Arthrobacter* is a predominant taxon in tobacco roots and becomes enriched in soils treated with nicotine. This bacterial genus catabolizes nicotine via the nicotine-catabolism (*nic*) gene cluster [29, 30] (Fig. S1). Comparative genome analyses demonstrated a prevalent distribution of the *nic* gene cluster in *Arthrobacter* strains derived from nicotine-containing environments, including tobacco roots and nicotine-treated soils. Recent studies have reported similar metabolic interactions, in which root microbiota catabolizes host-derived PSMs, in soybean (*Glycine max*), tomato, sesame (*Sesamum indicum*), and maize root microbiota [31–33]. These findings suggests that catabolic genes for host-specific PSMs are key to bacterial fitness in their host plant; however, whether these catabolic capacities directory contribute to host interactions and affect overall microbiota structure remains unclear.

Here, we investigated how bacterial catabolic capacities for host-derived PSMs shape both bacterial adaptation to host roots and root microbiota assembly, using the tobacco–*Arthrobacter* interaction system as a model. Comparative analysis of high-quality *Arthrobacter* genomes indicated that horizontal acquisition of the *nic* gene cluster is key for bacterial fitness in tobacco roots. To validate this genomic inference, we employed reverse genetics in both host and bacterial mutants impaired in nicotine biosynthesis and catabolism, respectively. Combined with gnotobiotic inoculation and SynComs experiments, we proposed a mechanistic model explaining the assembly process of plant species-specific root microbiota. This model suggests that the assembly is coordinated by the biosynthesis of host PSMs and their subsequent catabolism by the root microbiota.

## Methods

### Bacteria, plant, and chemical materials

The bacterial strains used in this study and their growth media are listed in Dataset S2. Except for NtRootC10, all strains were grown at 28°C in Luria–Bertani (LB) medium. NtRootC10 was grown at 28°C in TY medium (5 g/L tryptone, 3 g/L yeast extract) supplemented with 10 mM CaCl_2_. *Escherichia coli* DH5α were grown at 37°C in LB medium supplemented with appropriate antibiotics (50 μg/mL kanamycin). Seeds of *N. tabacum* cv. Petit Havana line SR-I were provided by Japan Tobacco, Inc. (Tokyo, Japan). Tobacco plants with the *erf189*/*erf199* mutant allele were generated using Petit Havana line SR-I as the genetic background in a previous study [34]. Chemicals were obtained from Wako Pure Chemical Industries (Japan) or Nacalai Tesque (Japan), unless otherwise indicated in Dataset S2.

### Isolation of root- and soil-derived bacteria strains

*Arthrobacter* strains were isolated from the soil (gray lowland soil) collected from a field at the Kyoto University of Advanced Science (KUAS), Kameoka, Kyoto, Japan (34°999380N, 135°559140E) with 9 independent experiments (Isolation1–9; DatasetS4). The detailed methods for Isolation 2–9 can be found in Shimasaki et al., 2021 and Nakayasu et al., 2021, respectively [19, 23]. In this study, we further isolated bacteria strains from tobacco roots grown in a growth chamber condition. Tobacco seeds (*Nicotiana tabacum* cv. Burley21) were surface sterilized with 70% ethanol (EtOH) for 1 min and 1% sodium hypochlorite (NaClO) for 10 min and rinsed 5 times with sterile water. Plants were germinated on Murashige and Skoog (MS) medium supplemented with 0.8% agar. Plants were grown in a growth chamber set at 25°C for 3 days in darkness and 7 days under long-day conditions (16 h light/8 h dark), transferred to plastic pots filled with soil from KUAS fertilized with Hyponex (Hyponex Japan, Japan), and grown for 5 weeks. After washing with tap water, roots were sectioned into smaller fragments and then homogenized with a mortar and pestle in PBS. Homogenates were diluted and plated onto LB agar, and incubated for up to 7 days at 28 °C. Single colonies were picked from plates, sub-cultured on LB agar, and preserved in a 25% glycerol solution at –80°C. Genomic DNA was extracted by the hot-alkaline DNA extraction method using extraction buffer (25 mM NaOH, 0.2 mM EDTA, 40mM Tris-HCl [pH 6.8]). The 16S *rRNA* genes were amplified using primer set 16S_10F and 16S_1500R (Dataset S3). PCR products were purified with the Wizard genomic DNA purification kit (Promega, Madison, USA) according to the manufacturer’s protocol and directly sequenced using primer 10F for taxonomic identification.

### Whole-genome sequencing and comparative genome analysis

Genomic DNA extraction, whole-genome sequencing, and assembly followed previously described procedures [23]. Comparative genome analysis incorporated publicly available *Arthrobacter* genome retrieved from the Integrated Microbial Genomes & Microbiomes system (https://img.jgi.doe.gov) and the NCBI (https://www.ncbi.nlm.nih.gov). The prediction and annotation of putative open reading frames (ORFs) of each genome was performed using Prokka [35]. High-resolution phylogenetic inference was conducted using Orthofinder [36]. The identification of homologous *nic* genes was performed by BLASTP using amino acid sequences of *Paenarthrobacter nicotinovorans* DSM420 against the proteome predicted by the *Arthrobacter* genome sequences (E-value thread > 1 × 10^−15^). Phylogenetic trees and associated data were visualized using ggtree [37].

### Plasmid construction and generation of gene knockout mutants

All constructs and primers used in this study are listed in Dataset S3. All insertion mutants of strain NtRootA2 were generated via homologous recombination. DNA fragments containing partial sequences of target genes were amplified via PCR using the genomic DNA of NtRootA2 as template and cloned into pCR4-TOPO vector (Thermo Fisher Scientific, USA) by TA cloning according to the manufacturer’s protocol. Transformation of NtRootA2 was performed by electroporation as described in Niu et al. [38]. For electrocompetent cell preparation, *Arthrobacter* sp. NtRootA2 was cultured in 100 mL of LB medium at 30°C until OD_600_ reached 0.3–0.5. Penicillin G was added to the medium at a final concentration of 30 μg/mL, and cultures were incubated for 1 h before harvesting via centrifugation (4°C, 4,000 × g, 10 min). Cells were washed three times with ice-cold electroporation buffer (0.5 M sorbitol containing 10% glycerol), resuspended, and stored at −80°C. Electrocompetent cells were mixed with 1 μg of plasmid DNA, and pulsed using a MicroPulser electroporator (Bio-Rad, USA) in a 2-mm cuvette at 2.5 kV, 600 Ω, and 10 μF. The bacterial cells were immediately transferred to 900μL of LB and incubated for 8 h at 30°C. The insertion mutants were selected on LB medium supplemented with 100 μg/mL kanamycin, and target gene disruptions were confirmed via PCR amplification.

### Conjugation experiment

To select transconjugants, a spontaneous rifampicin-resistant mutant of *Arthrobacter* sp. StoSoilB3 was generated. StoSoilB3 was grown overnight in LB at 28°C and subcultured in LB containing increasing concentrations of rifampicin (10, 25, 50, 100, 150 and 200 µg/mL). Stable rifampicin-resistant mutants were selected on LB agar plates containing 200 µg/mL rifampicin. The rifampicin-resistant mutant of StoSoilB3 and the VC mutant of NtRootA2 were grown in LB media supplemented with 200 µg/mL rifampicin or 100 µg/mL kanamycin, respectively. After three washes with sterile distilled water, each strain was resuspended in sterile distilled water at OD_600_ = 1.0 and mixed. The bacterial mixture was then spotted onto 0.22-μm MCE membranes (Merck Millipore, United States) placed on LB agar plates and incubated at 28°C overnight.

Transconjugants were selected on LB medium supplemented with 100 μg/mL of kanamycin and 200 μg/mL rifampicin. Horizontal plasmid transfer was confirmed via PCR and a nicotine degradation assay. Primers were designed to amplify the *ndhL* gene located on the pNtRootA2-2 plasmid and to the genomic regions unique to NtRootA2 and StoSoilB3 (Fig. S2 and Dataset S3).

### Measurements of nicotine-degrading activities using resting cells

Bacterial strains were cultured at 28°C in their respective media until reaching the stationary phase. Cells were collected by centrifugation at 13,000 × *g* for 2 min and washed twice with PBS buffer (137 mM NaCl, 1.8 mM KH_2_PO_4_, 10 Na_2_HPO_4_ •12H_2_O, 2.7mM KCl, pH 7.4). Pellets were resuspended in 2 mL PBS containing 1 mM nicotine at OD_600_ = 1.0. For SynCom degradation assays, equivalent amounts of each strain (OD_600_ = 1.0) were combined. After incubation for 2 days at 28°C, supernatants were collected by centrifugation at 13,000 × *g* for 2 min, filtered through a 0.45-μm Minisart RC4 filter (Sartorius, Germany), and stored at −30°C until ultra-high-performance liquid chromatography (UPLC) analysis.

The microbial reaction products were analyzed using an Acquity UPLC H-Class PLUS system equipped with a photodiode array (PDA) detector (Waters, United States) or a D-2000 Elite UPLC system with an L-2455 photodiode array detector (Hitachi, Japan). For HPLC analyses, 2 μL of each sample was injected onto an Atlantis Premier BEH C18 column (1.7 μm, 2.1 × 100 mm; Waters, United States) with a UPLC BEH C18 VanGuard Precolumn (1.7 μm, 2.1 × 5 mm^2^, Waters, United States). Mobile phases were 10 mM NH_4_HCO_3_ (pH 9.0; phase A) and acetonitrile (phase B). Elution was performed at 0.2 mL/min with 10% B isocratic for 0.25 min, followed by a linear gradient from 10% to 98% B over 6.75 min and 98% B isocratic for 2 min. For analyses on the D-2000 Elite HPLC System, 2 μL of each sample were injected onto an TSKgel ODS-80TM column (5.0 μm, 4.6 × 250 mm^2^; Toso, Japan). Elution was set at 0.2 mL/min with 40% B isocratic for 0.25 min, followed by a linear gradient from 40% to 98% B over 4.0 min and 98% B isocratic for more than 2 min. All analyses were conducted with the column oven temperature at 40°C, and UV absorbance at 264 nm was monitored using the PDA detector.

### In vitro bacterial growth assay

For growth assays using nicotine as the sole carbon source, the wild type, Δ*ndhL*, and VC mutants of NtRootA2 were grown in LB medium at 28°C. Cells were harvested via centrifugation at 13,000 × *g* for 1 min, washed with minimal medium (1×M9 minimal salt supplemented with 0.01% yeast extract, 0.01% MgSO_4_•7H_2_O, 0.005% CaCl_2_•2H_2_O, 0.00033% Fe(Ⅲ)-EDTA), and resuspended in minimal medium containing 5 mM nicotine at OD_600_ = 1.0. After overnight incubation at 28°C, cells were washed again, adjusted to OD_600_ = 0.1 in minimal medium containing 5 mM nicotine, and cultured at 28°C. For time-course bacterial growth analysis, 50 μL of culture supernatant was collected at 0, 2, 4, 8, 10, 12 and 16 h, centrifuged at 13,000 × *g* for 1 min, and resuspended in 50 μL of minimal medium. Growth of cultures was monitored based on OD_600_. Minimal medium supplemented with 5 mM glucose served as the control to confirm baseline growth capacity.

For growth assays in rich medium supplemented with nicotine, the wild type, Δ*ndhL*, and VC mutants were cultured in 2 ml TY medium at 28°C. Overnight cultures were washed with 10 mM MgCl_2_ and adjusted to OD_600_ = 0.001. then, 2 μl culture was added to 200 μl TY medium containing 1, 10, and 100 μM nicotine in a 96-well microplate. Growth curves were recorded at 600 nm every 15 min at 28°C for 24 h using a Varioskan LUX plate reader (Thermo Fisher Scientific, USA). For statistical analysis, the area under the curve was calculated using the bayestestR package [39].

### Bacterial colonization experiments

Bacterial strains were cultured at 28°C in LB medium to stationary phase, and then washed and resuspended in 10 mM MgCl_2_ at OD_600_ = 1.0. For competition assays, equivalent amounts of each bacterial strain were combined. Next, 8 μl of this suspension was added to 80 mL of 1/2 MS medium to obtain a final OD_600_ of 0.0001, then poured into plant boxes containing sterilized vermiculite. Sterilized tobacco seeds were germinated and grown on 1/2 × MS medium with 0.8% agar, and 10-day-old seedlings were transferred to plant boxes and grown for 20 days under long-day conditions (16 h light/8 h dark) at 25°C.

After 20 days of cultivation, the whole roots were washed with sterile water to remove surrounding vermiculite and placed in a 2-mL tube containing 1 mL 10 mM MgCl_2_ and 5–6 glass beads (2.3 mm). Samples were vortexed for 10 min to release root-colonizing bacteria, after which roots were transferred to a new 1.5-mL tube and weighed. The remaining suspension was diluted 1,000-fold and plated on LB agar plates and selection media. After 48 h of incubation at 28°C, colony-forming units (CFUs) were counted. For competitive colonization assays between wild-type NtRootA2 (Kan^S^) and the Δ*ndhL* or VC mutants (Kan^R^), wild-type CFUs were calculated by subtracting mutant CFUs (counted on LB medium containing 100 μg/mL kanamycin) from the total CFUs (counted on LB medium).

### SynCom inoculation and bacterial community analysis using 16S rRNA gene amplicon sequencing

All bacterial strains were cultured at 28°C in their respective media to stationary phase, collected by centrifugation at 13,000 × *g* for 2 min and washed twice with 10 mM MgCl_2_. Pellets were resuspended in 10 mM MgCl_2_ to OD_600_ = 1.0, and equal volumes of each strain were combined. Then, 8 μl of suspension was added to 80 mL of 1/2 MS medium (final OD_600_ of 0.0001) and poured into plant boxes filled with sterilized vermiculite. Sterilized tobacco seeds were germinated on 1/2× MS medium with 0.8% agar, and 10-day-old seedlings were transferred to plant boxes and grown for 20 days under long-day conditions (16 h light/8 h dark) at 25°C.

Collected tobacco roots were washed with sterile water to remove residual vermiculite, flash-cooled in liquid nitrogen, and ground at 3,000 rpm for 15 s using a Multi-Beads Shocker (Yasui Kikai Co., Japan). Fresh root weight for dPCR analysis was measured after removing residual vermiculite. Total DNA was extracted from the disrupted root tissues using the DNeasy PowerSoil Kit (Qiagen, Germany) according to the manufacturer’s protocol. Libraries were prepared using the two-step PCR amplification protocol of Kumaishi et al. [40]. The QIIME2 platform (version 2021.04) was used for sequence data analysis [41]. For all paired reads, the first 20 bases of both sequences were trimmed (to remove primer sequences), and the bases after 220 were truncated (to remove low-quality sequence data). Potential amplicon sequencing errors were corrected using DADA2 to produce an ASV dataset [42]. Obtained ASVs were aligned using MAFFT [43], and a phylogenetic tree was contracted using FastTree 2 [44]. Each ASV was assigned using a naïve Bayes classifier from the SILVA 138 rRNA database [45], and then the reads that were not assigned to SynCom members were removed. Obtained data was normalized by rarefying to 1,000 reads per samples for community analysis. The calculation of UniFrac distances was performed using the QIIME2 platform.

### Statistical analysis for microbiome profiling

All statistical analyses were performed in R software. Constrained principal coordinates analyses were performed based on weighted UniFrac distances using the capscale functions in the VEGAN package [46]. PERMANOVA (999 permutations) based on weighted UniFrac distances was performed to evaluate the effects of plant genotype and bacteria phenotype on community composition (model: beta diversity ~ plant genotype * SynCom phenotype).

### Absolute quantification of NtRootA2 colonization levels by digital PCR

The total copy number of the *socD* gene from strain NtRootA02 in the extracted DNA was quantified using the QIAcuity Digital PCR System (Qiagen, Germany). PCR amplification was performed in a 12-μl reaction mixture containing 1 μl template DNA, 4 μl QIAcuity EvaGreen PCR Master Mix (Qiagen, Germany), 0.6 μl each primers A02_socD_F and A02_socD_R primers (10 μM), and 5.8 μl RNase-free water. PCR conditions were 95°C for 2 min; 30 cycles of 95°C for 15 s, 62°C for 15 s, and 72°C for 15 s, followed by 40°C for 5 min. Copy number the *socD* per microliter of extract was corrected for elution volume and normalized to root fresh weight.

## Results

### The genus Arthrobacter has acquired the nic gene cluster via horizontal gene transfer

We previously demonstrated that the distribution of the *nic* gene cluster within the genus *Arthrobacter* is strongly associated with the presence of nicotine or related metabolites in their source environments [23]. To gain deeper insight into the ecological functions and evolutionary trajectory of this gene cluster, we isolated *Arthrobacter* strains from the roots of tobacco, soybean, and tomato and subjected them to whole-genome sequencing. We also isolated *Arthrobacter* strains from soils treated with different PSMs, including nicotine and santhopine from tobacco [23], and tomatine from tomato [19]. In total, 34 complete *Arthrobacter* genomes were obtained. By integrating publicly available sequences, we established an *Arthrobacter* pan-genome comprising 62 high-quality genomes (Dataset1). Although our *Arthrobacter* strains were isolated from the same soil, the *nic* gene cluster was specifically identified in the strains from nicotine-containing environments, such as tobacco roots or nicotine-treated soils (Fig. 1a). Within the entire *Arthrobacter* pan-genome, the *nic* gene cluster showed a sporadic distribution and predominantly found in isolates derived from nicotine or pesticide-containing environments (*P* = 2.97 × 10^−6^, Fisher’s exact test). Because several *Arthrobacter* strains carried the *nic* gene cluster on their plasmid (Figs. 1a and b), we inferred that these *Arthrobacter* strains may have acquired it via plasmid-mediated horizontal gene transfer (HGT) [47].

**Fig. 1.**
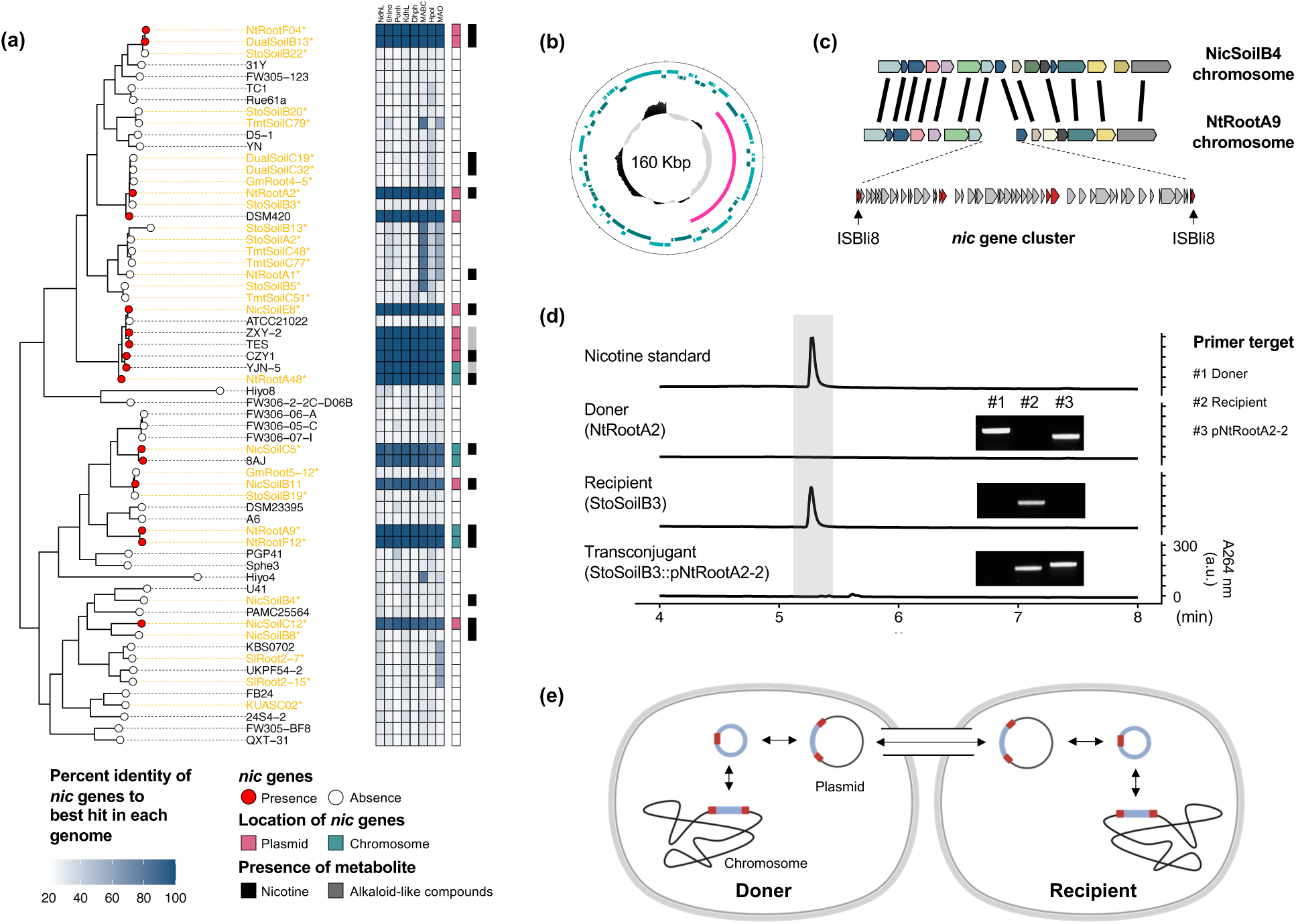
Genomic rearrangements in the genus *Arthrobacter* via horizontal transfer of the *nic* gene cluster. **(a)** Species tree of the genus *Arthrobacter* showing the phylogenetic distribution of the *nic* gene cluster. The phylogenetic tree of 62 *Arthrobacter* genomes was inferred from aligned single-copy genes using the MLE method. *Arthrobacter* strains isolated in this study are marked in orange with asterisk. Leaf nodes indicated presence or absence of the *nic* gene cluster. The heatmap showes percent identity of best BLASTP hits in each genome to the *nic* genes from *Paebarthrobacter nicotinovorans* pAO1 [48]. The inner bar indicates the location of the *nic* gene cluster. The outer bar shows the presence of nicotine or pesticides in the isolation environment. **(b)** Diagram of the *nic* gene cluster of strain NtRootA2 located on pNtRootA2-2 plasmid and **(c)** composite transposon carrying the *nic* gene cluster in strain NtRootA9. The composite transposon is flanked by IS elements ISBli8 (black arrows). This genome structure lies within the homologous genomic region of non-nicotine-degrading strain NicSoilB4. **(d)** Nicotine degradation assay and PCR amplifications of the donor, recipient, and transconjugant. Culture supernatants from strains grown in the nicotine-supplemented PBS buffer were analyzed via UPLC. All chromatograms were recorded at 264 nm using identical intensity scales. a.c., arbitrary units. Primer pairs amplified specific genomic regions of NtRootA2 and StoSoilB3, as well as the *ndhL* gene on pNtRootA2-2 plasmid. **(e)** Proposed model of genomic rearrangements in the genus *Arthrobacter* mediated by horizontal transfer of the *nic* gene cluster. The *nic* gene cluster can transfer to different genomic regions via a composite transposon within a single cell, and once integrated into conjugative plasmids, can transfer to other bacterial cells.

To test whether *Arthrobacter* can acquire the *nic* gene cluster via plasmid transfer, we performed a conjugation assay using a strain NtRootA2, which harbors the *nic* gene cluster on the pNtRootA2-2 plasmid (Fig. S1b). To trace their plasmid transfer, we inserted a kanamycin (Km)-resistance gene into a non-coding region of the pNtRootA2-2 plasmid, positioned outside the *nic* gene cluster (Fig. S1d). This modified donor strain was then co-cultivated with a recipient strain (StoSoilB3), which does not possess *nic* gene cluster, and transconjugants were selected by antibiotic selection (Fig. S2). Subsequent analysis confirmed the presence of pNtRootA2-2 in the transconjugant and demonstrated that it had gained nicotine degradation ability (Fig. 1d).

In contrast, some *Arthrobacter* strains harbor the *nic* gene cluster on their chromosome (Fig. 1a). Notably, *nic* gene clusters in strains NtRootA9 and F12 were flanked by insertion sequences (ISs), and the adjacent genomic region was strikingly conserved in the non-nicotine-degrading strain NicSoilB4 (Figs. 1c and S3). This arrangement—DNA located between two IS copies—constitutes a composite transposon, typically associated with intracellular DNA mobility [49]. Similar composite transposon-like structures were found on plasmid pZXY21 in strain ZXY-2 and plasmid pTES1 in strain TES (Fig. S3), implying that mobilization of the *nic* gene cluster by composite transposons is prevalent within the genus *Arthrobacter*. We therefore propose a transfer model in which the *nic* gene cluster moves between chromosomal and plasmid locations within a cell via composite and is then disseminated among bacterial cells when integrated into conjugative plasmids (Fig. 1e). Together, these findings suggest that HGT of the *nic* gene cluster mediated by mobile genetic elements facilitates *Arthrobacter* adaptation to nicotine-containing environments, including tobacco roots.

### Nicotine affects Arthrobacter growth in catabolism-dependent manner

To assess how the horizontal acquisition of the *nic* gene cluster contributes to *Arthrobacter* fitness in nicotine-containing environments, we constructed an insertional mutant of strain NtRootA2 by integrating vector pCR4-TOPO into the coding region of *ndhL* (Fig. S1c), which encodes the enzyme subunit catalyzing the first step of nicotine degradation (Figs. 2a and S1a). To control for vector insertion effects, we also used the kanamycin-resistant NtRootA2 line produced during the conjugation assay as a vector control (VC) in subsequent experiments (Fig. S1d). Nicotine degradation assays confirmed the loss of nicotine degradation ability in Δ*ndhL*, while the VC retained its catabolic capacity (Fig. 2b). Consistently, nicotine blue, an intermediate of nicotine degradation (Fig. 2a), was absent in Δ*ndhL* mutant, verifying that disruption of *ndhL* impaired nicotine catabolism.

**Fig. 2.**
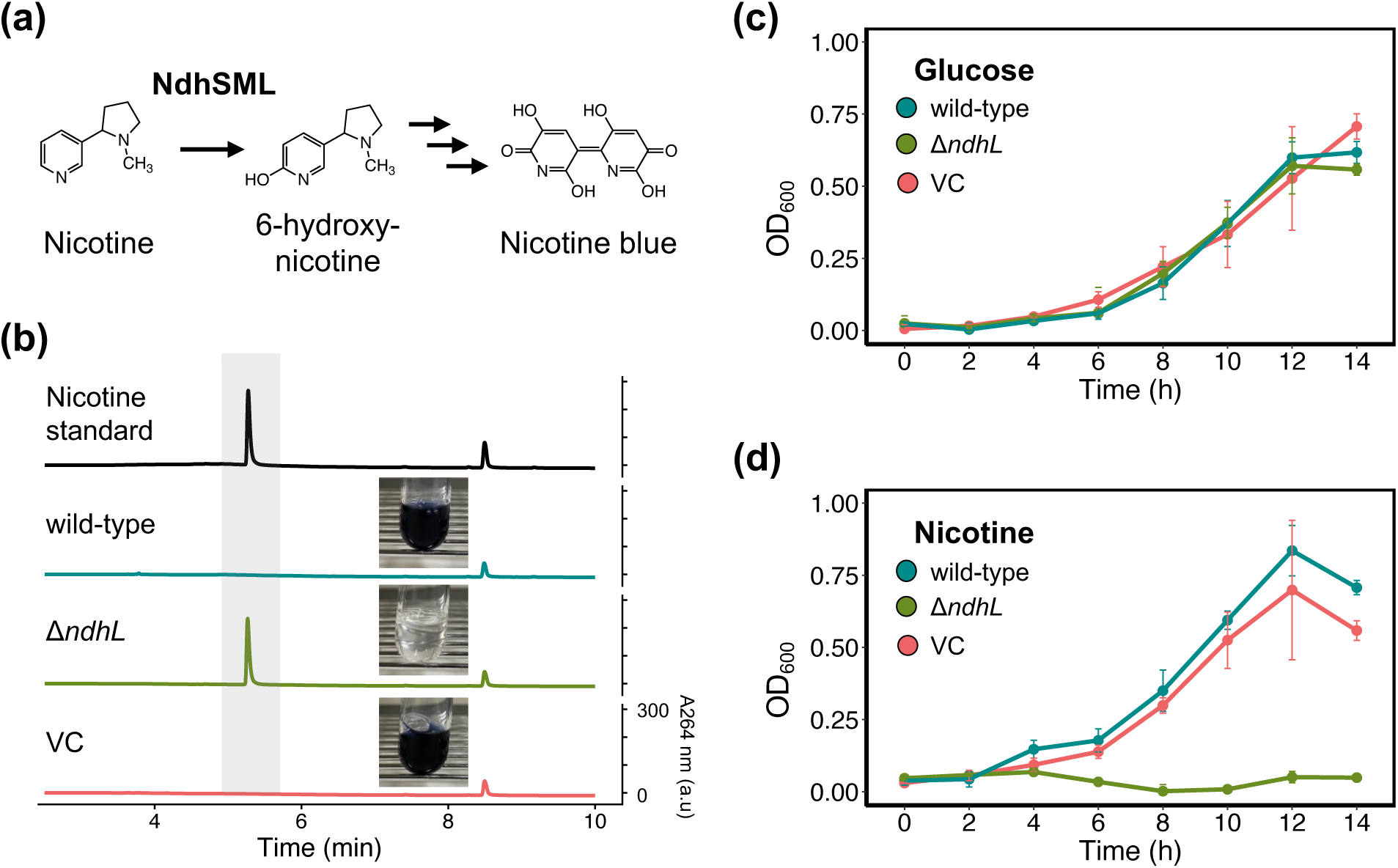
Nicotine affects *Arthrobacter* growth in a catabolism-dependent manner. **(a)** Partial nicotine degradation pathway in *Arthrobacter*. NdhSML hydroxylates the C6 position of pyridine ring in nicotine is to produce 6-hydroxynicotine. Bule-colored intermediates, known as nicotine blue, accumulates during degradation. **(b)** Nicotine degradation assay of NtRootA2 and its derivative mutants. UPLC analysis of culture supernatants grown in the nicotine-supplemented PBS is shown. All chromatograms were recorded at 264 nm using identical intensity scales. a.c., arbitrary units. Medium of wild-type and VC strains turned blue due to nicotine blue accumulation during nicotine catabolism. Growth rates of the wild-type, Δ*ndhL*, and VC strains of NtRootA2 in the minimal medium supplemented with **(c)** glucose or **(d)** nicotine are shown. Error bars indicate Mean ± SD of three replicates.

We cultured these strains in minimal medium supplemented with either glucose or nicotine as the sole carbon source. All strains grew to a similar level in glucose, but only Δ*ndhL* failed to grow in nicotine medium (Figs. 2c and d), indicating that NtRootA2 utilizes nicotine as a carbon source via the *nic* gene cluster. Beyond nutrient utilization, recent studies demonstrated that PSMs inhibit bacterial growth, while bacterial catabolism of these compounds increases bacterial tolerance by detoxifying them [33]. We therefore evaluated *in vitro* bacterial growth of NtRootA2 in nutrient media containing three different nicotine concentrations. Unexpectedly, the growth of NtRootA2 was not affected even in the presence of 100 μM of nicotine (Fig. S4). Importantly, nicotine also did not influence the growth of Δ*ndhL*, implying that *Arthrobacter* may exhibit nicotine tolerance, although this tolerance is independent with its catabolic capacity. While chemotaxis is another crucial bacterial trait involved in bacterial root colonization [50–52], canonical flagellar motility and chemotaxis genes were rarely conserved in NtRootA2 (Fig. S5). These results indicate that nicotine primarily affects *Arthrobacter* growth by serving as a nutrient source.

### Nicotine catabolism enhances competitive fitness of Arthrobacter in tobacco roots

*In vitro* growth assays showed that nicotine affects *Arthrobacter* growth in a catabolism-dependent manner. Next, we examined root colonization by the wild-type and mutant strains to determine the impact of nicotine-catabolic capacity on *Arthrobacter* adaptation to tobacco roots. In order to clarify the contributions of host-derive nicotine, we also employed a double-knockout mutant of jasmonate (JA)-responsive ETHYLENE RESPONSE FACTOR (ERF) transcription factor genes *ERF199* and ERF189 (*erf189*/*erf199*) [34], which produces significantly less amount of nicotine in leaves and roots. In wild-type tobacco roots, Δ*ndhL* colonized at significantly lower levels than the wild type and VC (Fig. 3a). However, in the *erf189*/*erf199* roots, all strains were colonized at comparable levels. Two-way analysis of variance (ANOVA) demonstrated significant effects of both *Arthrobacter* catabolism and plant nicotine biosynthesis on *Arthrobacter* colonization, with a significant interaction. Notably, host nicotine biosynthesis had a greater influence on *Arthrobacter* colonization than bacterial catabolism (*R^2^* Plant = 0.26, *R^2^*Bacteria = 0.06).

**Fig. 3.**
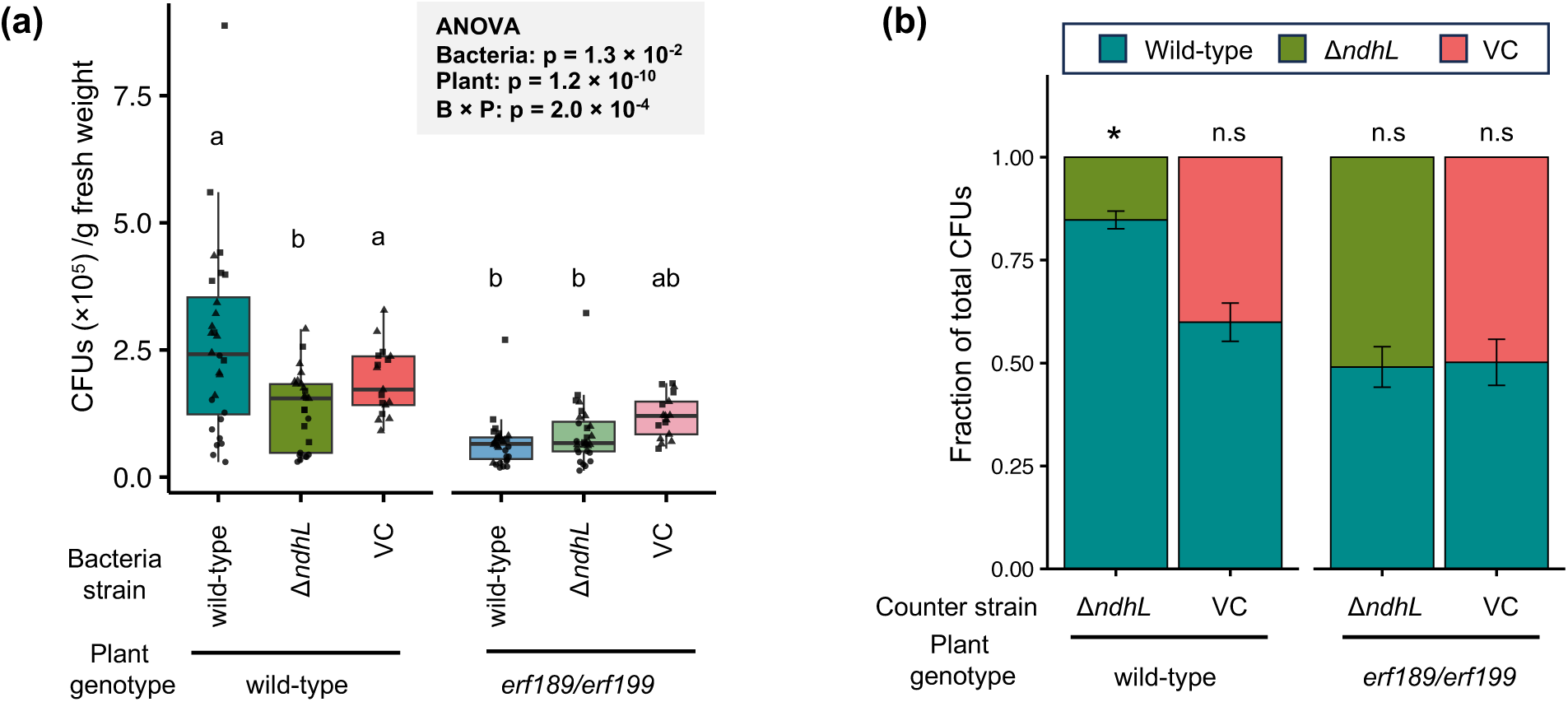
Nicotine-catabolic capacity contributes to *Arthrobacter* fitness in the tobacco roots. **(a)** Colonization assay of NtRootA2 and its derivative mutants in a mono-association system. These *Arthrobacter* strains were inoculated into the wild-type or *erf189*/*erf199* tobacco roots. Data are from three independent experiments, except for two groups (VC in the wild-type roots and VC in the *erf189*/*erf199* roots), which were performed twice (n = 17–30). Different letters indicate statistical differences corresponding to two-way ANOVA with Tukey’s honest significant difference (HSD) test (α = 0.05). Different shapes depict data points from full-factorial biological replicates. Bacteria: Bacteria strain (wild-type, Δ*ndhL*, or VC). Plant: tobacco genotype (Wild-type or *erf189*/*erf199*). “B × P”: interaction between bacteria phenotype and tobacco genotype. **(b)** Root colonization competition assay between the wild-type and Δ*ndhL* or VC strains in the roots of wild-type or *erf189*/*erf199* mutant tobacco. Data are from two independent experiments. Asterisks indicated statistical significance based on paired-samples *t*-tests before transformation to relative values for visualization (FDR-correlated q-value < 0.05). Bars indicate mean SD (n = 20–22). n.s., not significant.

We then investigated the role of nicotine catabolism under the competitive conditions within the root niche by co-inoculating the wild-type and eitherΔ*ndhL* or VC strains into wild-type or *erf189*/*erf199* mutant tobacco roots. In the wild-type tobacco roots, Δ*ndhL* showed significantly reduced colonization relative to the wild-type, whereas the VC colonized at a comparable level (Fig. 3b). No competitive differences were observed in both pairs of strains inoculated into the *erf189*/*erf199* roots. Together, these results indicate that nicotine catabolism enhances *Arthrobacter* fitness in tobacco roots, particularly under competitive conditions.

### Nicotine biosynthesis and its bacterial catabolism coordinately determine tobacco root microbiota community

Because *Arthrobacter* catabolism and tobacco biosynthesis of nicotine significantly affected *Arthrobacter* colonization of the tobacco roots, we next investigated whether these metabolic interactions also affect tobacco root microbiota assembly. To address this, we established SynComs comprising 11 phylogenetically diverse bacterial strains isolated from tobacco roots, including the wild-type strain of *Arthrobacter* sp. NtRootA2 (hereafter the wildSC) (Fig. 4a). Two additional SynComs were generated by substituting this wild-type strain either the Δ*ndhL* or VC strains (ndhLSC and conSC, respectively). Except for the NtRootA2 wild-type and VC strains, none of the SynCom members degraded nicotine (Fig. S6), whereby nicotine-catabolism abilities of *Arthrobacter* strains represented the entire nicotine-catabolism capacity of each SynCom (Fig. 4b). Based on this, the SynComs were classified as “Degrading SynComs” (wildSC and conSC) or “Non-degrading SynCom” (ndhLSC).

**Fig. 4.**
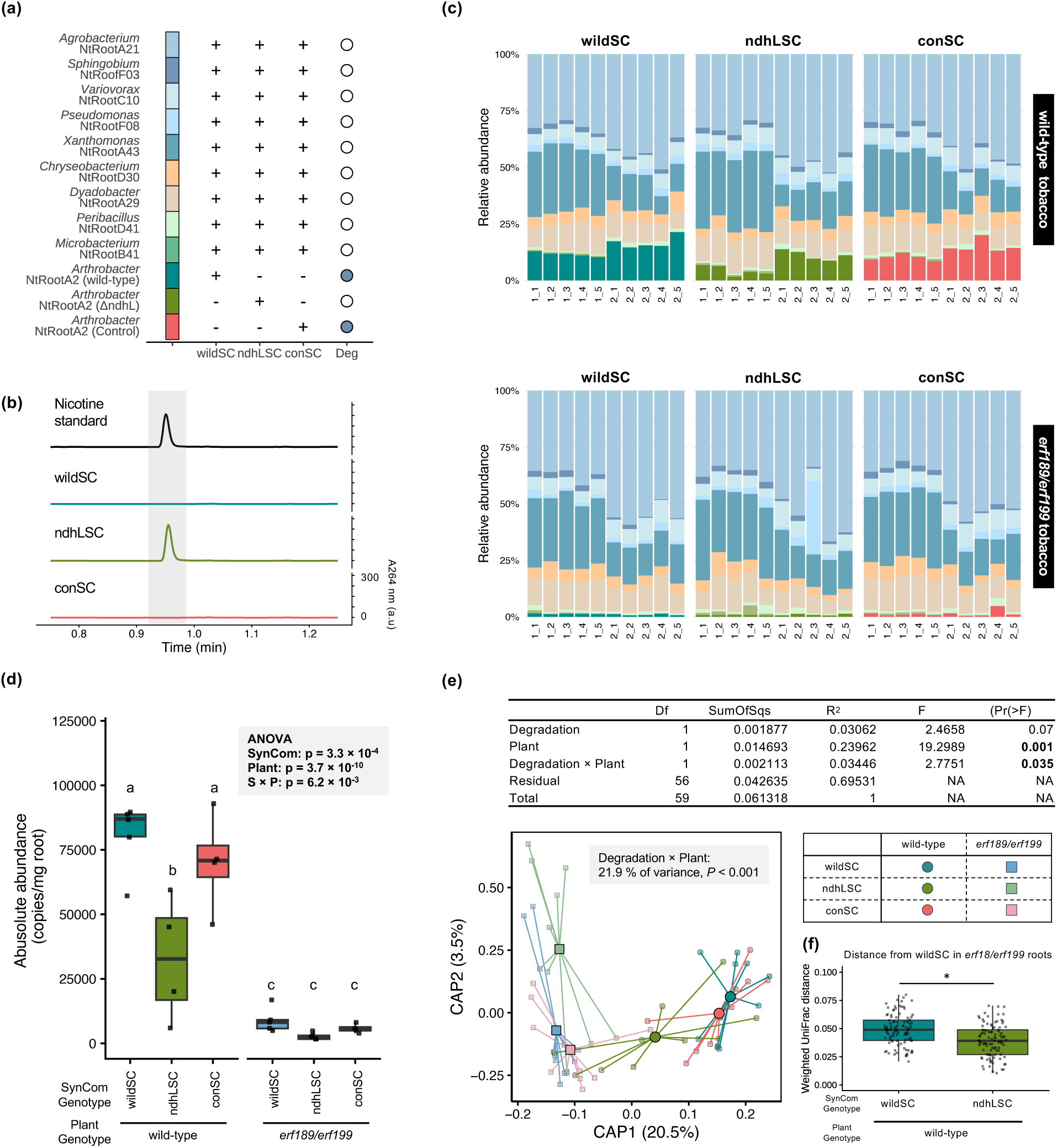
Nicotine biosynthesis and its bacterial catabolism jointly influence the community structure of the tobacco root microbiota. **(a)** Bacterial strains used in the synthetic community (SynCom). Strains included in each SynCom are maeked “+” and excluded strains “−”. Detailed taxonomic information is shown in Dataset S2. The right column indicates the nicotine degradation ability of each bacterial strain. Deg, nicotine degradation. **(b)** Nicotine degradation assay of each SynCom. HPLC analysis of culture supernatants from SynCom grown in nicotine-supplemented PBS is shown. All chromatograms were recorded at 264 nm using identical intensity scales. a.c., arbitrary units. Essentially identical results were obtained from two independent experiments. **(c)** Bacterial community composition of each SynCom inoculated into the wild-type or *erf189*/*erf199* tobacco roots. **(d)** Absolute abundance of strain NtRootA2. Individual data points are indicated with dots (n = 4–5). The result table for two-way ANOVA is included. SynCom: wildSC, ndhLSC, or conSC. Plant: tobacco genotype (Wild-type or *erf189*/*erf199*). S × P: interaction between SynCom and tobacco genotype. Different letters indicate statistical differences corresponding to two-way ANOVA followed with HSD test (α = 0.05). **(e)** Constrained principal coordinates analysis of the community structure of SymComs in wild-type and *erf189*/*erf199* roots based on weighted Unifrac distance. Data are from two independent experiments. Ordination was constrained by the model ~ Degradation phenotype × tobacco genotype. Degradation: SynCom phenotype (“Degrading SynCom” [wildSC, conSC] or “Non-degrading SynCom” [ndhLSC]). Plant: tobacco genotype (Wild-type or *erf189*/*erf199*). Degradation × Plant: interaction between SynCom phenotype and tobacco genotype. PERMANOVA results based on weighted UniFrac distance for variation in bacterial community structure are shown in the upper table. **(f)** Comparison of weighted UniFrac distance. Box plots showing mean and variance of average pairwise weighted UniFrac distances between samples of wildSC inoculated into *erf189*/*erf199* tobacco roots and samples of wildSC- or ndhLSC-inoculated into wild-type tobacco roots (n = 100). Asterisks indicated statistical significance (*t*-test, *P* < 0.05).

These SynComs were inculcated into wild-type or *erf189*/*erf199* tobacco roots and community structures were assessed via 16S *rRNA* gene amplicon sequencing. All strains except *Paenibacillus* NtRootD41 were detected in roots at 20 days after inoculation. *Agrobacterium* NtRootA21 and *Xanthomonas* NtRootA43 were the most dominant, followed by *Dyadobacter* NtRootA29, across both plant genotypes (Figs. 4c and S7). The relative abundance of NtRootA2 was significantly reduced in the ndlLSC in wild-type roots and in all SynComs inoculated into the *erf189*/*erf199* roots. Absolute quantification using digital PCR targeting *socD*, a single-copy gene unique to NtRootA2 showed a similar colonization pattern, confirming that these changes are quantitative (Fig. 4d). As in mono-association colonization assays (Fig. 2a), plant genotype excreted a stronger effect on NtRootA2 absolute abundance than SynCom phenotype (*R^2^* Plant = 0.67, *R^2^* SynCom Phenotype = 0.12), with a significant interaction between those factors. The relative abundance of *Variovorax* NtRootC10 and *Agrobacterium* NtRootA21 were significantly higher in the *erf189*/*erf199* root (*P* = 0.0325 and *P* = 0.0189, respectively) (Fig. S7), but were rarely influenced by the SynCom phenotype, suggesting that nicotine directly affects colonization of certain SynCom members. We also observed an interaction between plant genotype and SynCom phenotype on the relative abundance of *Dyadobacter* NtRootA29 (*P* = 0.0292), suggesting that nicotine mediates bacteria–bacteria interactions through its catabolism.

Permutational analysis of variance (PERMANOVA) revealed significant shifts in overall bacterial community composition driven by plant genotype (*R^2^*= 0.24, *P* = 0.001), whereas SynCom phenotype showed no major effect (*R^2^*= 0.03, *P* = 0.07) (Fig. 4e). Interestingly, despite minor effect of SynCom phenotype alone on the bacterial community structure, we detected a significant interaction between SynCom phenotype and plant genotype (*R^2^* = 0.03, *P* = 0.035). Notably, the disruption of *ndhL* shifted community composition toward that formed in the *erf189*/*erf199* roots inoculated with wildSC or conSC (Fig. 4f), which was largely explained by the first axis of the constrained ordination analysis (Fig. 4e). A slight community shift was also detected in the ndhLSC, even in *erf189*/*erf199* roots, likely reflecting the incomplete loss of nicotine biosynthesis in *erf189*/*erf199*, which still accumulates detectable nicotine levels in the roots [34]. Consistent with this, digital PCR showed a slight, but statistically non-significant reduction in the Δ*ndhL* mutant compared with the wild-type and VC in the *erf189*/*erf199* roots (Fig. 4d). Collectively, our results show that plant-derived nicotine biosynthesis and its bacterial catabolism jointly determine tobacco root microbiota structure.

## Discussions

### The function of PMSs in the assembly of plant species-specific root microbiota

Root exudates contain numerous primary and specialized metabolites that strongly influence root microbiota composition. Here, we provide experimental evidence that PSMs shape the root microbiota by serving as nutrient sources for specific root microbiota member, thereby conferring a competitive advantage (Fig. 4). Catabolism of host-derived PSMs by root microbiota has also been reported in various plant–bacterial systems [23, 31, 33], indicating that the ability to metabolize PSMs is a common adaptation strategy. Likewise, it is widely appreciated that primary metabolites play a fundamental role in microbial recruitment, as root-associated bacteria generally catabolized amino acids, organic acids, and sugars, more efficiently than non-adapted isolates [53–55]. Large-scale comparative genome studies further revealed that plant-associated bacteria harbor more genes related to carbohydrate metabolism functions compared to non-plant-associated bacteria [56, 57]. Importantly, these primary-metabolic traits are broadly conserved across among core root microbiota members. In contrast, PSM catabolic genes are generally restricted to a narrow subset of bacterial taxa [29, 33, 58], and exogenous application of PSMs to soil often enriches those capable of degrading them [59]. Together with our findings, these observations support the two-step selection process whereby PSMs act as host genotype factors that fine-tune root microbiota composition by selectively enriching bacterial taxa with the capacity to metabolize them.

### MGEs play a role in metabolic adaptation of root microbiota

Although we isolated *Arthrobacter* strains from the same soil, the *nic* gene cluster was identified only in strains from tobacco roots or nicotine-treated soils (Fig. 1a). Similarly, the isoflavone catabolism (*ifc*) and benzoxazinoid degradation (*bxd*) gene clusters are confined to root microbiota of legumes and maize, respectively, which produce these PSMs [33, 58]. These findings suggest that PSM catabolic genes are selectively distributed, even among strains within the same bacterial taxa. Our comparative genome study revealed that plasmid-mediated HGT explains this restricted genomic adaptation. In contrast, recently identified PSM catabolic genes such as *ifc* and *bxd* genes reside on the chromosome. This inconsistency may be explained by the larger size of the *nic* gene cluster, which comprises over 40 genes spanning approximately 50 kb, compared with the smaller *ifc* (~20 genes) and *bxd* (~15 genes) clusters. Since plasmid carriage imposes replication and segregation costs that can reduce bacteria fitness [60, 61], they are selectively maintained under a specific environment where they can confer strong benefits to the host bacteria. It is conceivable that a trade-off between the plasmid carriage cost and the metabolic advantage conferred by the *nic* gene cluster likely explains its restricted phylogenic distribution within *Arthrobacter*.

Transfers of MGEs have also been reported in other commensal bacteria, and these genomic rearrangement events appear to be facilitated by metabolites present in root exudates [62–64]. For instance, fabricated ecosystems showed conjugative plasmid transfer among *Pseudomonas putida* in the *Brachypodium distachyon* rhizosphere [65], with frequencies increasing in response to host-derived metabolites. Moreover, a long-read metagenomic analysis also uncovered numerous previously unknown plasmids in the rice leaf microbiota [66], highlighting the widespread but underexplored functions of MGEs in the plant–microbiota interactions. Therefore, given our results showing strong effects of HGT on *Arthrobacter* fitness and on tobacco root microbiota composition, functional characterizations of MGEs will further elucidate molecular mechanisms of bacterial host adaptation and root microbiota assembly.

### Multi-functional aspects of PSMs in root microbiota regulation

Despite the significant contribution of bacterial catabolism of host-derived PSMs in adaptation, the host nicotine-biosynthesis mutant exerted a stronger impact on bacterial colonization than the nicotine-catabolism mutant of *Arthrobacter* (Fig. 4d). This difference likely reflects the multi-functionality of PSMs in interactions with the root microbiota. Beyond serving as a nutrient source for *Arthrobacter*, nicotine also exhibits antimicrobial activity against various microbes [67]. We found that *Variovorax* NtRootC10 and *Agrobacterium* NtRootA21 were enriched in the *erf189*/*erf199* roots, whereas their abundances were not significantly influenced by the nicotine-catabolic capacity of the inoculated SynComs (Fig. S7). Given the minor effect of nicotine catabolism on their colonization, their enrichment may be driven by the less toxic environment in the *erf189*/*erf199* roots, enabling these strains outcompete *Arthrobacter*. Conversely, nicotine also stimulates growth of root-associated bacteria, such as *Pseudomonas aeruginosa* NXHG29 [28] and many PSMs act as molecular attractants, inducing chemotaxis and biofilm formation [50–52, 68]. Our findings, together with these studies highlight that PSMs influence bacterial behaviors through both catabolism-dependent and independent mechanisms, and these combined functions may shape the taxonomic and functional profiles of root microbiota.

Beyond adaptation to their host roots, degradation of host-derived PSMs also plays a role in bacteria–bacteria interactions. For example, degradation of isothiocyanates, a characteristic class of PSMs found in *Brassicaceae* plants, influences the growth of non-degrading commensals through cross-feeding of metabolic intermediates or detoxification [69, 70]. It has also been demonstrated that the presence of other bacterial strains can redirect the BX catabolic pathway of BXs of a degrading strain, likely by metabolizing intermediate compounds [71]. Nicotine is similarly metabolized by various bacterial species via at least three distinct pathways [29], producing metabolic intermediates such as nicotine blue and methylamine. Therefore, it is crucial to comprehensively understand how nicotine and its degradation intermediates are metabolized by the whole bacterial community and how these metabolic interactions shape root microbiota.

### Ecological relevance of catabolism-mediated root microbiota assembly

A major remaining question concerns the ecological relevance of nicotine catabolism-mediated assembly of the tobacco root microbiota. It has been well documented that biotic and abiotic stresses often stimulate PSM biosynthesis and secretion [72–74], suggesting that plants may secrete PSMs to establish the beneficial root microbiota that can alleviate these stresses [22, 75, 76]. This PSM-triggered functioning of root microbiota is referred to as a “cry for help” or “microbiome feedbacks” [77, 78]. In this study, we propose that bacterial catabolic capacities for host-derived PSMs contribute to these processes. For instance, cucurbitacins, the characteristic bitter triterpenoids in cucurbit plants, recruit *Enterobacter* and *Bacillus*, thereby enhancing resistance to soil-borne wilt fungal pathogen [20]. Notably, this feedback effect of the *Enterobacter* was mediated by cucurbitacin degradation. *Arthrobacter* colonization can promote plant growth and modulate plant hormone signaling pathways, including auxin and JA signaling [79–81]. Hence, it is intriguing to investigate whether the nicotine catabolism-driven community shift in tobacco roots participates a “microbiome feedbacks” response to environmental stresses, such as herbivore attacks, which trigger nicotine biosynthesis [25].

## Conclusion

Root microbiota can catabolize various compounds secreted by plant roots, and these catabolic capacities mediate complex plant–microbiota interactions [82]. In this study, we show that the nicotine-catabolic capacity of *Arthrobacter* substantially contributes to its colonization of tobacco roots. Comparative genome analysis demonstrates that mobile gene elements (MGEs) underlie this metabolic interaction between tobacco and *Arthrobacter*. These results support a mechanistic model, in which horizontal acquisition of the *nic* gene cluster improves *Arthrobacter* fitness and confers a competitive advantage over other microbiota members, thereby shaping the characteristic tobacco root microbiota (Fig. 5). This study advances our molecular understanding of how PSMs shape plant-species-specific root microbiota.

**Fig. 5.**
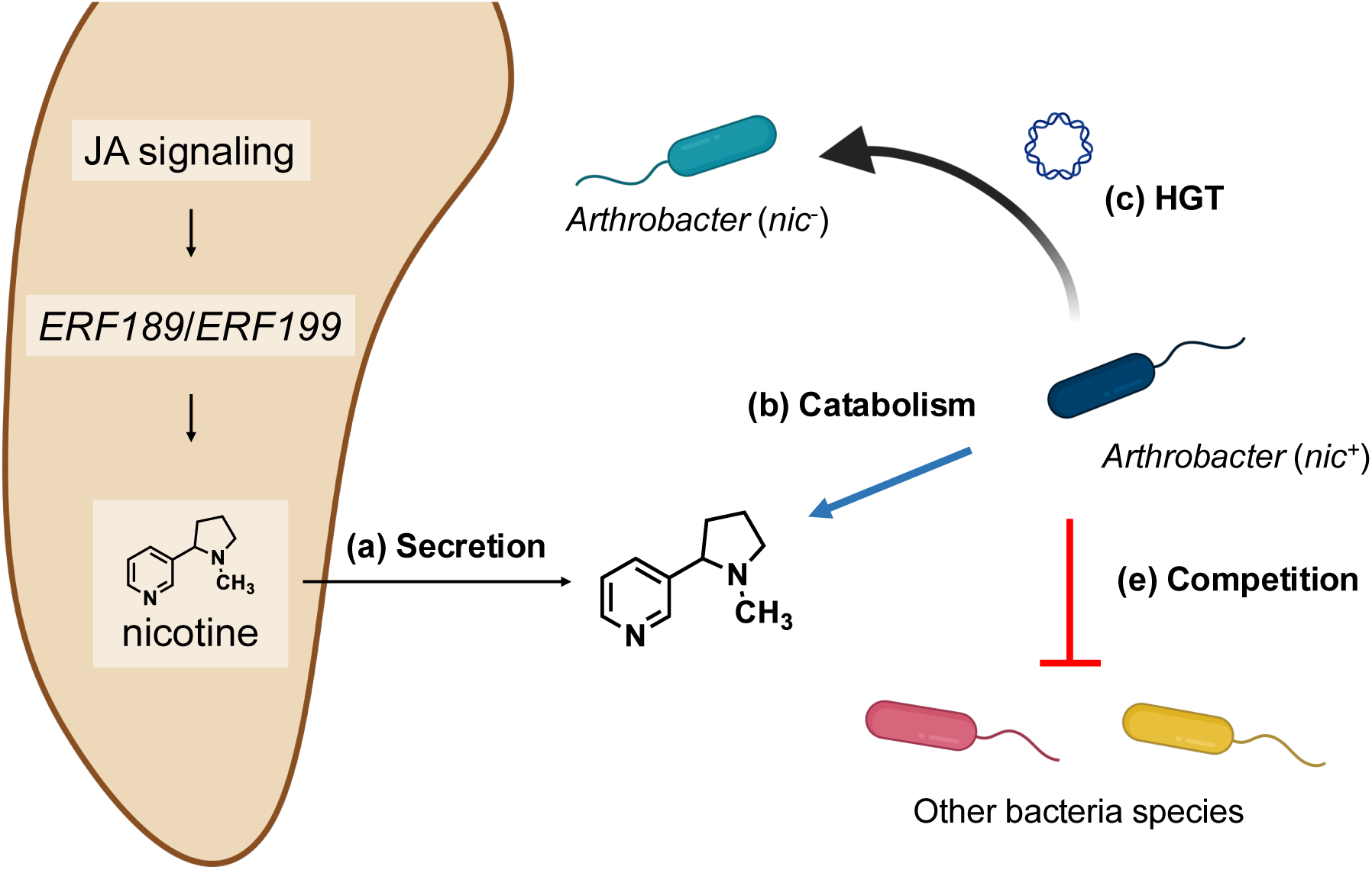
Proposed model of nicotine-mediated assembly of the tobacco root microbiota. Anticipated events related to the interaction between tobacco and its-associated microbiota are represented by letters. **(a)** Nicotine secretion promotes the formation of tobacco-specific root microbiota characterized by the enrichment of *Arthrobacter*. **(b)** The *nic* gene cluster enables this bacterial adaptation to tobacco roots and **(c)** this gene cluster can horizontally transfer among the genus *Arthrobacter* via plasmid transfer. **(d)** Nicotine-catabolic capacity confers a competitive advantage on *Arthrobacter* over other microbiota members, shaping a characteristic tobacco root microbiota. The figure was created with Biorender.com.

## Declarations

### Approval and Consent to participate

Not applicable.

### Consent for publication

Not applicable.

### Availability of data and materials

The data set supporting the results of this study is publicly available at the DDBJ (https://www.ddbj.nig.ac.jp) (PRJDB11299 and PRJDB18225). Additional data related to this paper will be made available from the corresponding author upon reasonable request.

### Competing interests

The authors have no conflicts of interest directly relevant to the content of this article.

### Founding

This study was supported in part by Japan Society for the Promotion of Science (JSPS) Research Fellowship for Young Scientists PD (22KJ3147 to T.S.); and JSPS KAKENHI grants (22K21367 to R.T.N.); and the Mayekawa Houonkai Foundation to T.S; and the Humanosphere Science Research of RISH, Kyoto University, to T.S, and RIKEN TRIP initiative to K.S.

### Author’s contributions

Tomohisa Shimasaki, Yasunori Ichihashi, Akifumi Sugiyama, and Ryohei Thomas Nakano conceived and designed the research. Tsubasa Shoji, Ken Shirasu, Kazufumi Yazaki, Yasunori Ichihashi, Akifumi Sugiyama, and Ryohei Thomas Nakano supervised the experiments. Tomohisa Shimasaki, Sachiko Masuda, Arisa Shibata, and Ken Shirasu conducted the whole-genome sequencing. Tomohisa Shimasaki and Ryohei Thomas Nakano analysed bacterial genome data. Tomohisa Shimasaki generated the *Arthrobacter* mutants and conducted bacterial fitness assays both *in vitro* and *in planta*. Tomohisa Shimasaki, Maiko Furubayashi, and Yoshitomo Kikuch measured nicotine degradation ability. Tomohisa Shimasaki, Yui Nose, Shuhei Yabe, and Yasunori Ichihashi performed the SymCom inoculation assays and analyzed the bacterial community data. Tomohisa Shimasaki and Ryohei Thomas Nakano wrote the manuscript with contributions from all the authors. Tomohisa Shimasaki and Ryohei Thomas Nakano agree to serve as the authors responsible for contact and ensure communication.

## Acknowledgments

We thank Ms. Mine Tabuse, Ms. Yoshiko Shimizum, and Ms. Akiko Tsuruta for technical assistance; We thank Dr. Hiroaki Adachi and Dr. Tadashi Fujiwara for providing access to the Varioskan LUX, which greatly facilitated this study. We would like to thank Dr. Masaru Nakayasu and Dr. Ryosuke Munakata for helpful discussion; Japan Tobacco, Inc., for providing the tobacco seeds.

## Supplemental figure captions

**Fig. S1. Nicotine-catabolism pathway and *nic* gene cluster of *Arthrobacter* spp., and generation of NtRootA2 mutants. (a)** Overviwe of the nicotine degradation by *Arthrobacter* spp. and **(b)** and *nic* gene cluster of NtRootA2. The cluster comprises three sub-gene clusters involved in γ-N-methylaminobutyrate catabolism, nicotine degradation, and molybdenum cofactor biosynthesis, represented in purple, blue, and green, respectively. Insertion sequences are shown in red. **(c)** An internal fragment of the *ndhL* gene was cloned into the pCR4-TOPO vector to generate the *ΔndhL* mutant. **(d)** For the VC mutant, the genomic region overlapping loci 04174–04175, located outside the *nic* gene cluster, was targeted. Both Δ*ndhL* and VC mutant of NtRootA2 were generated via a single crossover-mediated homologues recombination.

**Fig. S2. Conjugation experiment for the horizontal transmission of the *nic* gene cluster.** The donor strain NtRootA2 carried a plasmid with a kanamycin resistance gene. A spontaneous rifampicin-resistant mutant of strain StoSoilB3, which lacks the *nic* gene cluster, served as the recipient. Black triangles indicate the primer sets targeting doner- and recipient-specific genomic regions and the *ndhL* gene. Donor (Kan^R^/Rif^S^) and recipient (Kan^S^/Rif^R^) strains were co-cultured on LB medium. Transconjugants (Kan^R^/Rif^R^) were selected on LB medium containing 100 μg/mL kanamycin and 100 μg/mL rifampicin. Transmission of the *nic* gene cluster was confirmed by PCR amplification and a nicotine degradation assay.

**Fig. S3. The *nic* gene clusters of *Arthrobacter* strains used for comparative genome analysis.** The three sub-clusters of the *nic* gene cluster (γ-N-methylaminobutyrate catabolism, nicotine degradation, and molybdenum cofactor biosynthesis) are shown in purple, blue, and green, respectively. Insertion sequences are shown in red. IS elements flanking the *nic* gene cluster are indicated by white-filled arrows.

**Fig. S4. *In vitro* growth assay of NtRootA2 and its derivative mutants in TY medium supplemented with three different concentrations of nicotine. (a)** Growth curves (OD_600_) of each *Variovorax* isolates in TY medium supplemented with different nicotine concentrations. Data are expressed as mean ± SD (n = 5). Control: 0 μM nicotine; Low, 1 μM nicotine; Medium, 10 μM of nicotine; and High, 100 μM nicotine. **(b)** The area of the growth curve (AUC) was calculated based on OD_600_ measures over 24 h. Data points were analyzed based on one-way ANOVA with Dunnett’s post hoc test between control and nicotine. n.s., not significant. Data are presented as mean ± standard deviation with individual data points (n = 5).

**Fig. S5. KEGG pathway maps for flagellar assembly (ko02040) and bacterial chemotaxis (map02030) in strain NtRootA2.** Genes detected in the genome of *Arthrobacter* sp. NtRootA2 are highlighted in green.

**Fig. S6. Nicotine degradation assay of SynCom members.** UPLC analysis of culture supernatants from each bacteria strain grown in nicotine-supplemented PBS. Chromatograms were recorded at 264 nm using identical intensity scales. a.c., arbitrary units. Essentially identical results were obtained from two independent experiments.

**Fig. S7. Mean relative abundance of single strains in the SynComs.** Means ± SE, boxplots, and individual datapoints are shown (n = 10). Data were obtained from two independent experiments. The two-way ANOVA result table is provided. Different letters indicate statistical differences based on two-way ANOVA followed by HSD test (α = 0.05). SynCom: wildSC, ndhLSC, or conSC. Plant: tobacco genotype (Wild-type or *erf189*/*erf199*). S × P: interaction between SynCom and tobacco genotype. Different letters indicate statistical differences corresponding to two-way ANOVA followed by HSD test (α = 0.05).

## Notes

### Competing Interest Statement

The authors have declared no competing interest.

