## Supplemental figures for "Horizontal acquisition of nicotine catabolism gene cluster drives the assembly of tobacco root microbiota community"

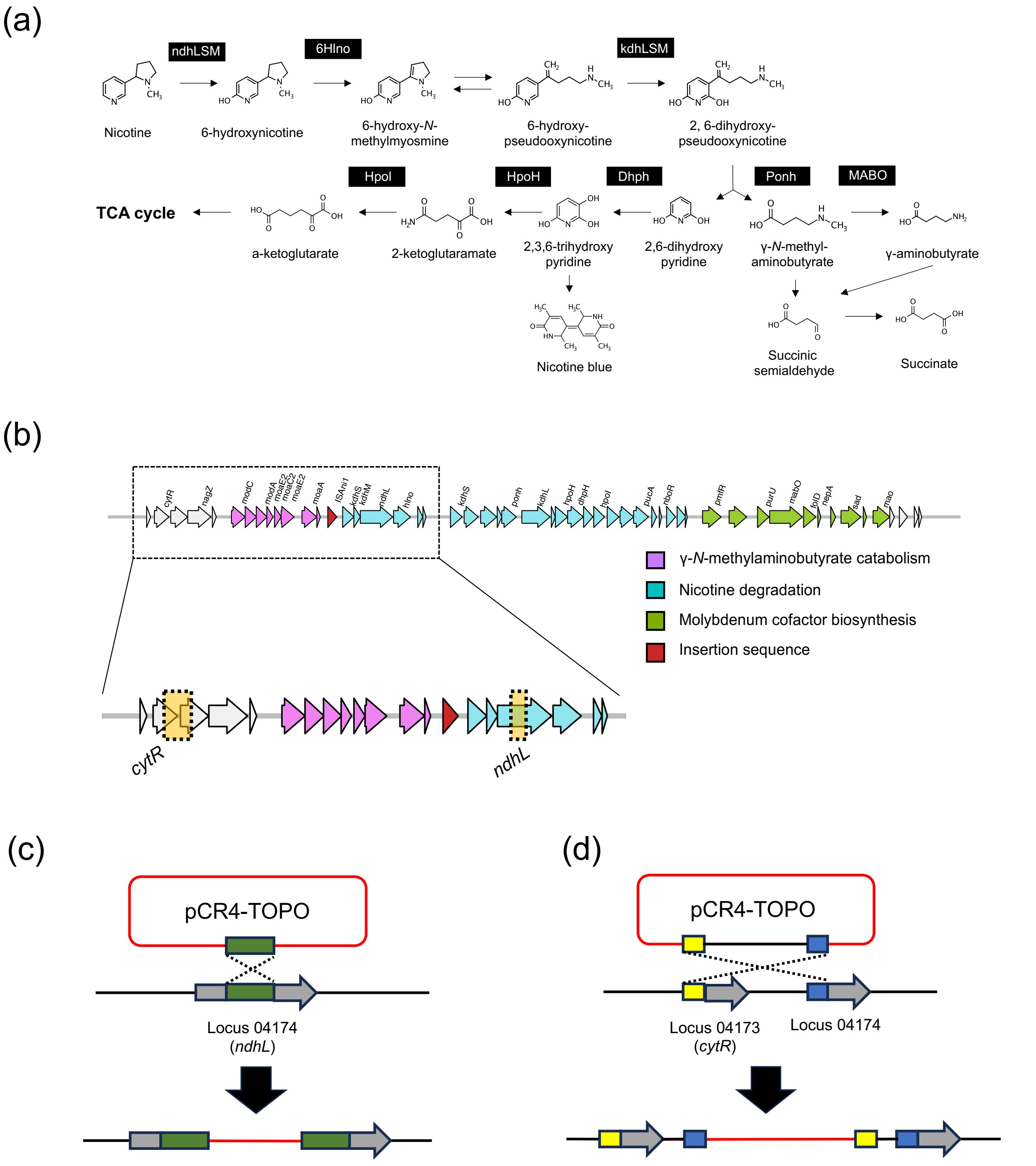


**Fig. S1. Nicotine-catabolism pathway and *nic* gene cluster of *Arthrobacter* spp., and generation of NtRootA2 mutants. (a)** Overviwe of the nicotine degradation by *Arthrobacter* spp. and **(b)** and *nic* gene cluster of NtRootA2. The cluster comprises three sub-gene clusters involved in γ-N-methylaminobutyrate catabolism, nicotine degradation, and molybdenum cofactor biosynthesis, represented in purple, blue, and green, respectively. Insertion sequences are shown in red. **(c)** An internal fragment of the *ndhL* gene was cloned into the pCR4-TOPO vector to generate the *ΔndhL* mutant. **(d)** For the VC mutant, the genomic region overlapping loci 04174–04175, located outside the *nic* gene cluster, was targeted. Both Δ*ndhL* and VC mutant of NtRootA2 were generated via a single crossover-mediated homologues recombination.

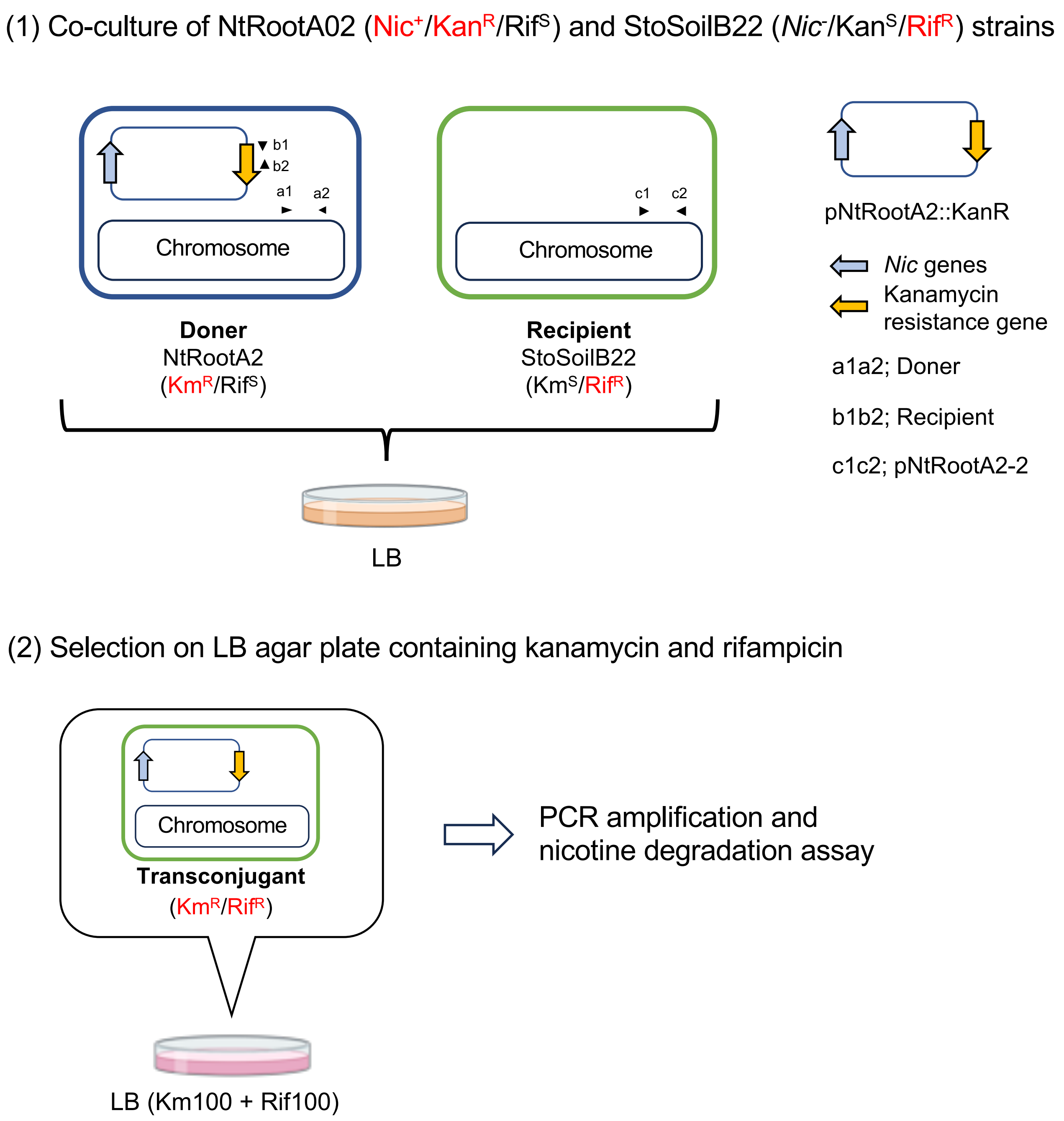


**Fig. S2. Conjugation experiment for the horizontal transmission of the *nic* gene cluster.** The donor strain NtRootA2 carried a plasmid with a kanamycin resistance gene. A spontaneous rifampicin-resistant mutant of strain StoSoilB3, which lacks the *nic* gene cluster, served as the recipient. Black triangles indicate the primer sets targeting doner- and recipient-specific genomic regions and the *ndhL* gene. Donor ($Kan^R$/$Rif^S$) and recipient ($Kan^S$/$Rif^R$) strains were co-cultured on LB medium. Transconjugants ($Kan^R$/$Rif^R$) were selected on LB medium containing 100 μg/mL kanamycin and 100 μg/mL rifampicin. Transmission of the *nic* gene cluster was confirmed by PCR amplification and a nicotine degradation assay.

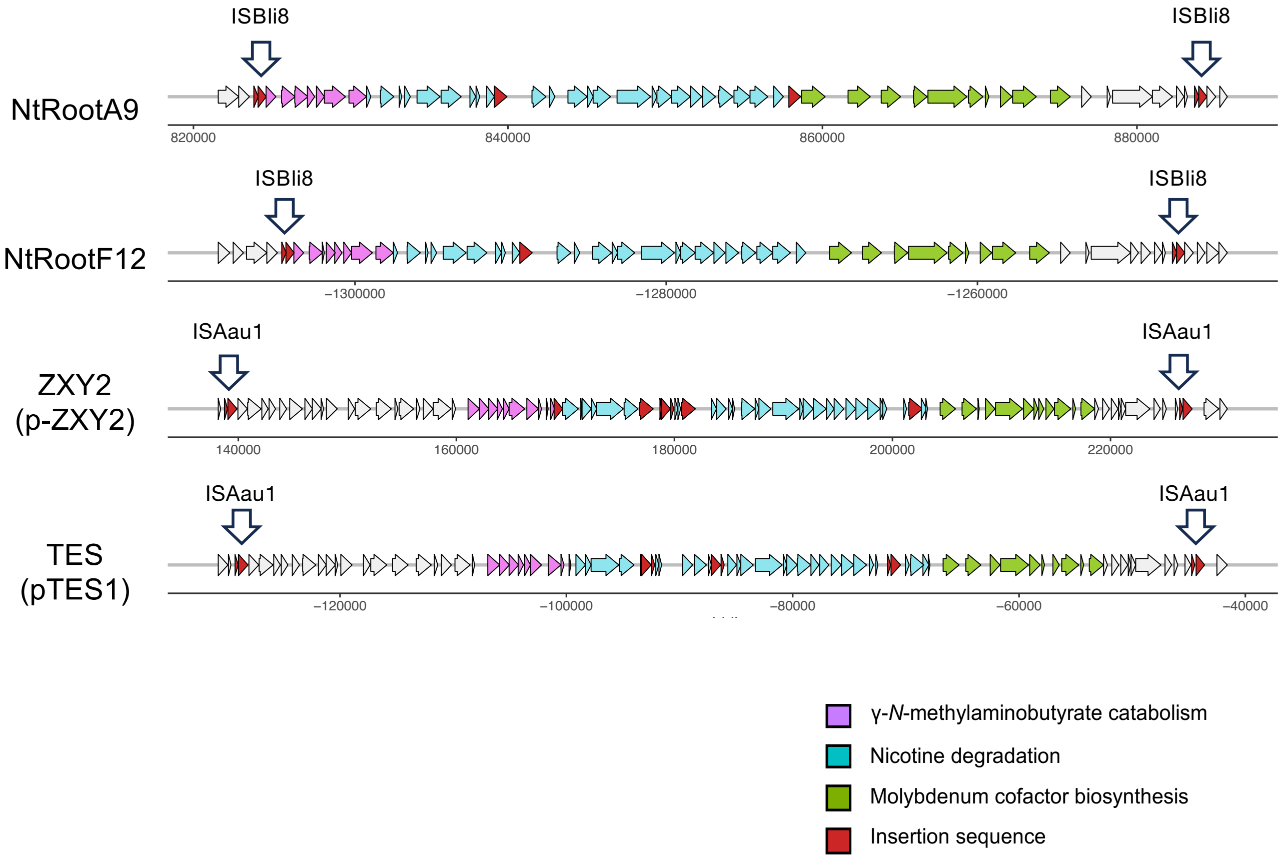


**Fig. S3. The *nic* gene clusters of *Arthrobacter* strains used for comparative genome analysis.** The three sub-clusters of the *nic* gene cluster (γ-N-methylaminobutyrate catabolism, nicotine degradation, and molybdenum cofactor biosynthesis) are shown in purple, blue, and green, respectively. Insertion sequences are shown in red. IS elements flanking the *nic* gene cluster are indicated by white-filled arrows.

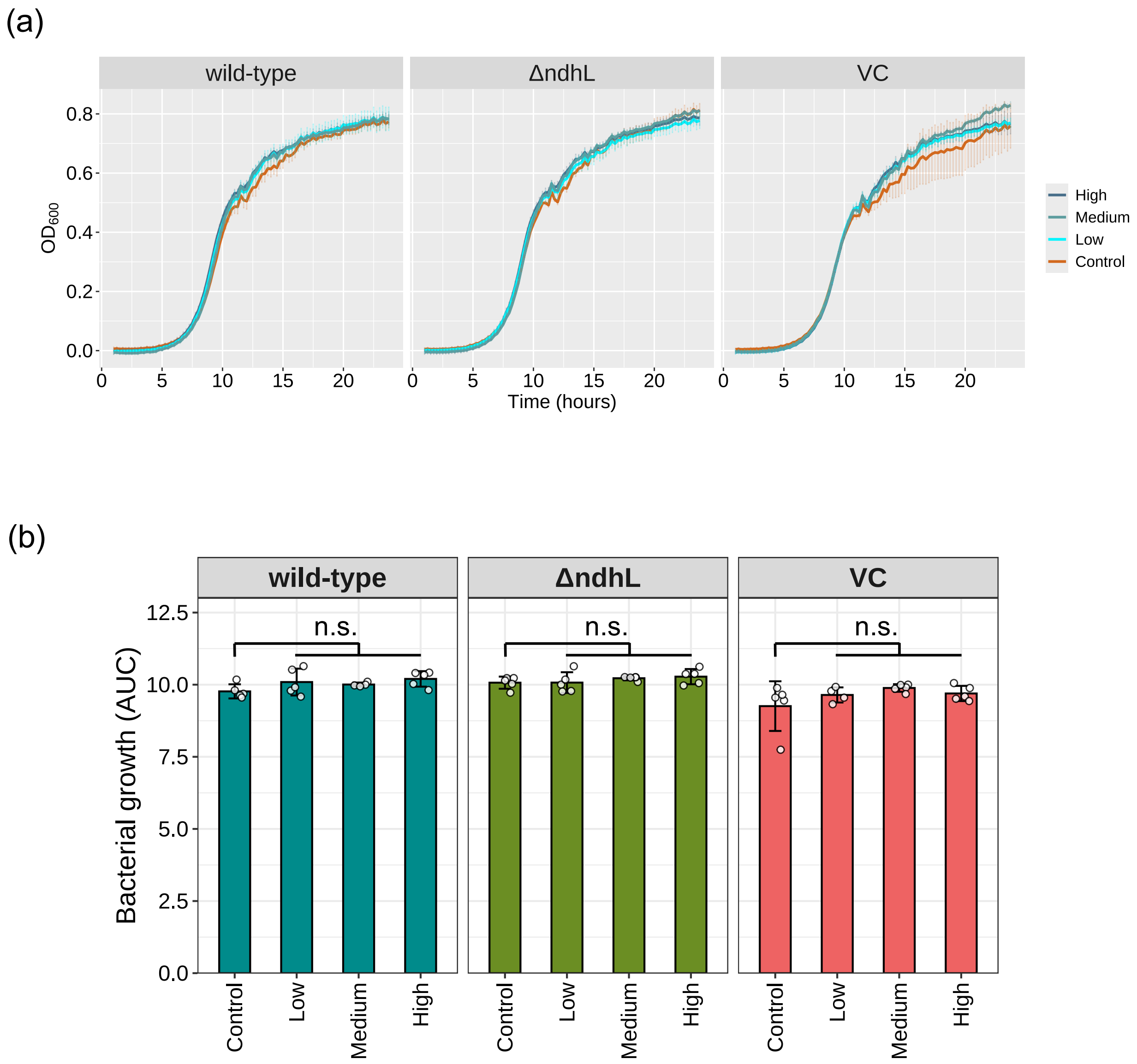


**Fig. S4. *In vitro* growth assay of NtRootA2 and its derivative mutants in TY medium supplemented with three different concentrations of nicotine. (a)** Growth curves ($OD_{600}$) of each *Variovorax* isolates in TY medium supplemented with different nicotine concentrations**.** Data are expressed as mean ± SD (n = 5). Control: 0 μM nicotine; Low, 1 μM nicotine; Medium, 10 μM of nicotine; and High, 100 μM nicotine. (b) The area of the growth curve (AUC) was calculated based on $OD_{600}$ measures over 24 h. Data points were analyzed based on one-way ANOVA with Dunnett's post hoc test between control and nicotine. n.s., not significant. Data are presented as mean ± standard deviation with individual data points (n = 5).

(a)

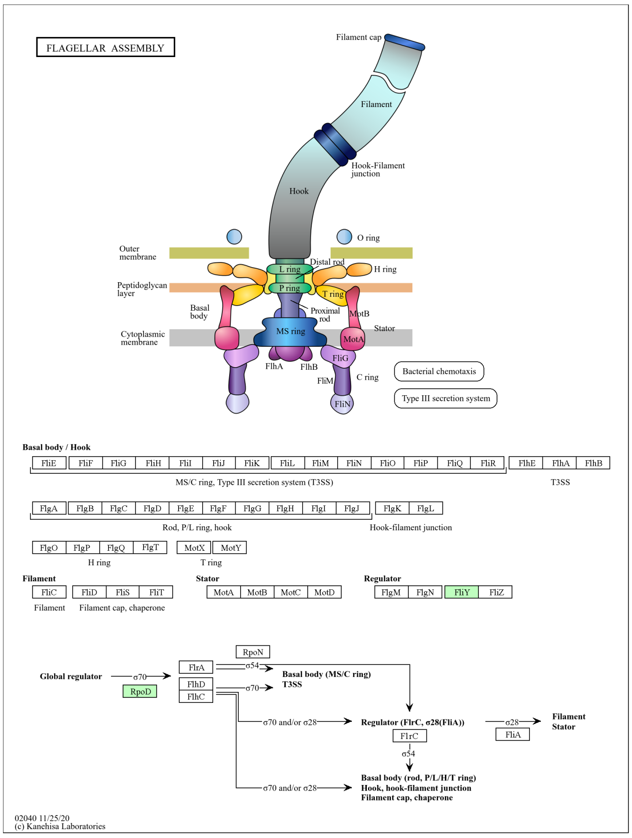


(b)

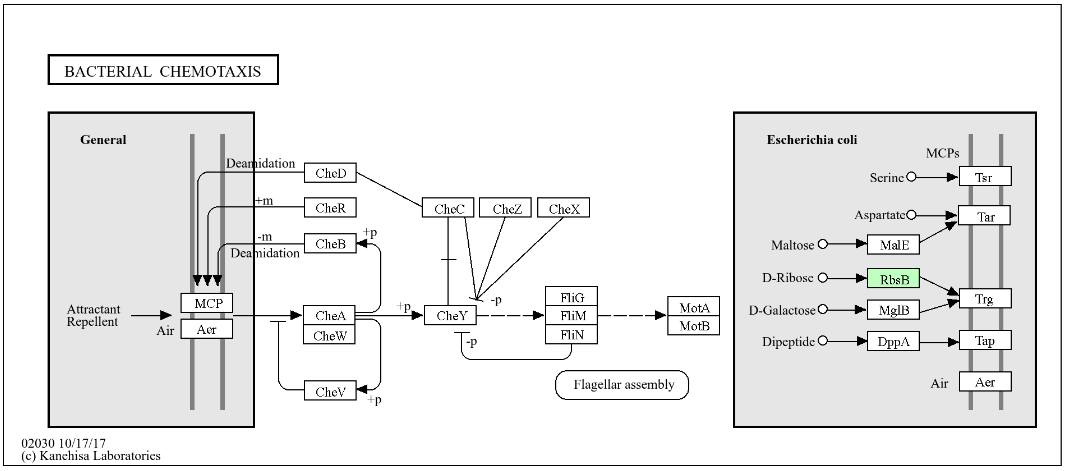


**Fig. S5. KEGG pathway maps for flagellar assembly (ko02040) and bacterial chemotaxis (map02030) in strain NtRootA2.** Genes detected in the genome of *Arthrobacter* sp. NtRootA2 are highlighted in green.

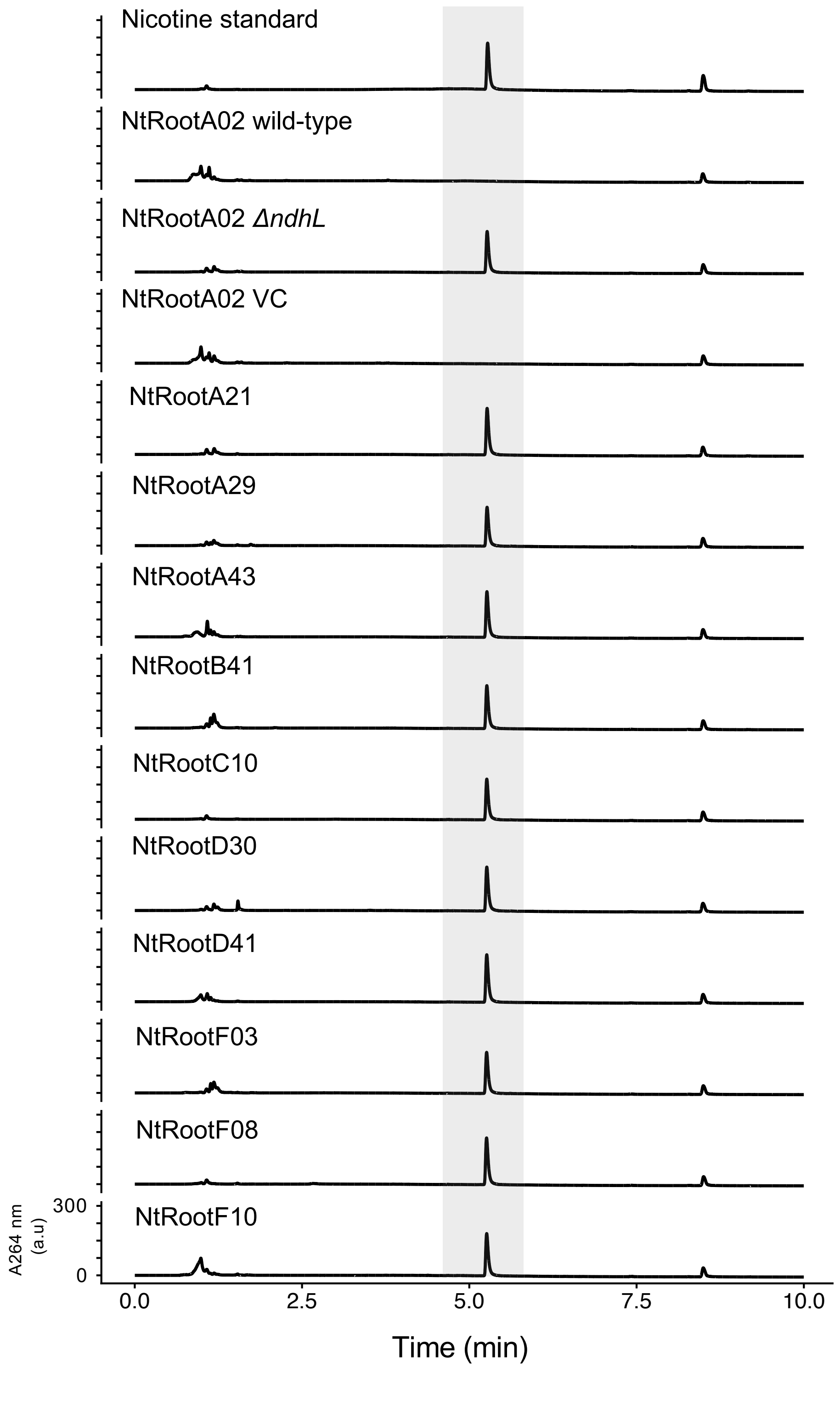


**Fig. S6. Nicotine degradation assay of SynCom members.** UPLC analysis of culture supernatants from each bacteria strain grown in nicotine-supplemented PBS. Chromatograms were recorded at 264 nm using identical intensity scales. a.c., arbitrary units. Essentially identical results were obtained from two independent experiments.

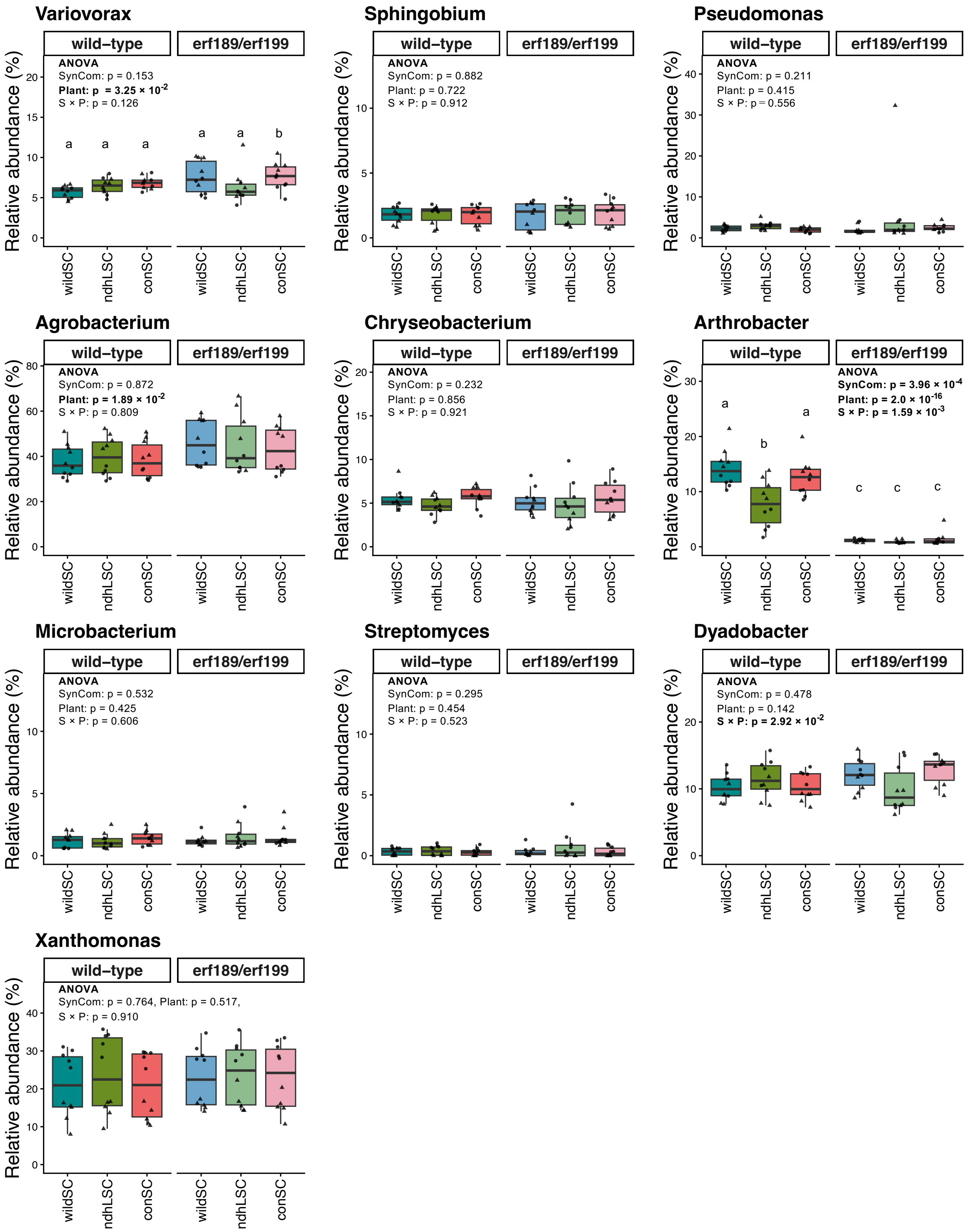


**Fig. S7. Mean relative abundance of single strains in the SynComs.** Means ± SE, boxplots, and individual datapoints are shown (n = 10). Data were obtained from two independent experiments. The two-way ANOVA result table is provided. Different letters indicate statistical differences based on two-way ANOVA followed by HSD test (α = 0.05). SynCom: wildSC, ndhLSC, or conSC. Plant: tobacco genotype (Wild-type or *erf189/erf199*). S × P: interaction between SynCom and tobacco genotype. Different letters indicate statistical differences corresponding to two-way ANOVA followed by HSD test (α = 0.05).
